# A putative genomic map for resistance of *Bos indicus* cattle in Cameroon to bovine tuberculosis

**DOI:** 10.1101/2020.04.26.057497

**Authors:** Rebecca Callaby, Robert Kelly, Stella Mazeri, Franklyn Egbe, Lindert Benedictus, Emily Clark, Andrea Doeschl-Wilson, Barend Bronsvoort, Mazdak Salavati, Adrian Muwonge

**Author notes:** Correspondence: Rebecca Callaby.

## Abstract

Bovine Tuberculosis (bTB) caused by *Mycobacterium bovis* is a livestock disease of global economic and public health importance. There are currently no effective vaccines available for livestock and so control relies on animal level surveillance and pasteurisation of dairy products. A new alternative control approach is to exploit the genetic variability of the host; recent studies have demonstrated that breeding *Bos taurus* cattle for increased resistance to bTB is feasible. The utility of such an approach is still unknown for the *Bos indicus* cattle population. This study aims to assess genetic variation in bTB resistance and the underlying genomic architecture in *Bos indicus* breeds in Cameroon.

We conducted a cross-sectional study of slaughter cattle in Cameroon and genotyped a sample of 213 cattle. Their genomic diversity was characterised using PCA, hierarchical clustering and admixture analysis. We assessed genetic variation in bTB resistance using heritability analysis and compared quantitative trait loci.

Previous studies had found that breed was an important factor in explaining the epidemiology of bTB, with Fulani cattle appearing to be more susceptible than mixed breeds. However, we show that the apparent phenotypic differences in visual appearance between the breeds was not reflected by clear genomic differences. At the genetic level, cattle belonging to different hierarchical genomic clusters differed in their susceptibility to bTB. There was evidence of a genomic association between *M. bovis* infection status with specific SNPs.

We highlight the need to understand the challenges faced by livestock in specific settings both in terms of pathogens and the environment, in addition to their intended purpose and how they fit into a defined management system. It is only at this point livestock keepers can then make informed breeding choices, not only for resistance to disease but also for increasing production.

## 1 INTRODUCTION

Bovine tuberculosis (bTB) caused by *Mycobacterium bovis* is a major zoonotic livestock disease causing a chronic respiratory condition characterised by weight loss and poor welfare and eventually death. In low-and middle-income countries (LMICs) it is estimated that M. *bovis* is responsible for approximately 1.4% of human TB cases, which equates to an estimated 70,000 new human infections annually in Africa (Olea-Popelka et al., 2017). *Mycobacterium bovis* is a member of the *Mycobacterium tuberculosis* complex, it primarily infects humans via consumption of unpasteurised milk, poorly cooked meat and close contact with infected animals. Unfortunately, the risks of bTB infection from milk are poorly understood by the livestock keepers in Cameroon, who are mainly pastoralist (Kelly et al., 2016; Ayele et al., 2004). There are currently no effective vaccines available for use in livestock and public health control relies on pasteurisation of dairy products, surveillance at post-mortem examination in slaughterhouses and active surveillance of cattle as part of test and slaughter programmes. LMICs in sub-Saharan Africa currently employ passive abattoir surveillance through official veterinary services, while pasteurisation is carried out at the household level in many settings where there is no centralised collection of milk. In these settings, bTB is considered endemic, with the prevalence in cattle estimated to range from 6-20% depending on the region and diagnostic tools used (Dibaba and Daborn, 2019; Asante-Poku et al., 2014).

*M. bovis* has a wide host range, which includes domestic and wild bovids, small ruminants, swine and cervids; which means that control requires a strong coordinated multi-disciplinary, trans-boundary approach. Unfortunately, most LMICs have poorly resourced veterinary services and do not have the means to implement bTB control strategies, instead they rely on controlling the risk of exposure through abattoir surveillance. At the same time LMICs, encouraged and funded by many charities, have widely adopted dairy development strategies with the aim of aiding livelihood and community development (Bill & Melinda Gates Foundation, 2012; Heifer International, 2019; Tambi, 1991). Although this brings clear nutritional and economic benefits to those keeping dairy cattle, there are concerns that the current strategy based on the introduction of exotic (European taurine) genetic characteristics i.e. improved milk production and faster growth in the local breeds of Africa, may also result in increased susceptibility to and risk of several zoonotic diseases including bTB (Opoola et al., 2019; Opoola, 2018; Hanotte et al., 2000; Bahbahani et al., 2018; Coffie et al., 2015).

There is epidemiological evidence from Ethiopia that there are breed differences in susceptibility to *M. bovis* with *Bos indicus* breeds appearing to be less susceptible compared to European Holsteins *Bos taurus* (Vordermeier et al., 2012). Breed introductions are usually based on a centripetal model i.e. governments import taurine breeds into centrally located breeding facilities, which then distribute “improved breeds” outwards into rural areas (Mekonnen et al., 2019). In Ethiopia, for example, concerns have been raised that this approach could inadvertently spread bTB from urban areas where the rates are high to rural settings where they are lower (Mekonnen et al., 2019).

To mitigate such risks new approaches to add to the current bTB control tool kit are urgently required. One such approach is breeding for resistance to bTB. This is based on exploiting the observed genetic variation in resistance to *M. bovis* infection in cattle (Tsairidou et al., 2014; Brotherstone et al., 2010; Bermingham et al., 2014). Furthermore, recent research demonstrated that breeding for increased resistance to bTB is feasible, and has generated the necessary tool set to carry out selective breeding for bTB resistance (Tsairidou et al., 2018; Banos et al., 2017). The UK dairy industry has recently implemented pedigree selection into the “TB Advantage” genetic evaluation index that enables breeders and farmers to select bulls with greater bTB resistance (Banos et al., 2017). This has proven to be popular with farmers and the breeding industry.

There is, therefore, an opportunity to help prevent or at least reduce the scale of an emerging bTB epidemic in African dairy cattle through genetic selection. For this approach to gain traction, we need to reconcile the potential disparity between phenotypic and genotypic characteristics of cattle in Africa. Our previously published results (Kelly et al., 2018; Kelly, 2017) show that even after controlling for other factors, breed (within the *Bos indicus* sub species) is an important factor in explaining the increased risk of infection, with the Fulani breed appearing to be more susceptible (Kelly et al., 2018; Kelly, 2017). This raises the tantalising possibility of local phenotypic breeds (as characterised by the local abattoir employees) being more resistant to bTB.

In this paper we investigate this possibility by using genetic and phenotypic data from cattle diagnosed as bTB positive and negative during a cross-sectional study of bTB in Cameroon (Egbe et al., 2016, 2017; Kelly et al., 2018) to assess genetic variation in bTB resistance and the underlying genomic architecture in Bos *indicus* breeds in Cameroon.

## 2 MATERIALS AND METHODS

### 2.1 Study area and sampling

A cross-sectional study of 2346 slaughter cattle was conducted in four regions of Cameroon, see Figure 1. As animals came into each abattoir the age, as estimated by the dentition score (individuals were defined as young if they have a dentition score between 0-2 i.e. no permanent incisors; old individuals have a dentition score between 3-5), sex and breed as defined by the local abattoir employees (mixed breed or Fulani *Bos indicus*) was recorded. A heparinized blood sample was collected and local Ministry of Livestock, Fisheries and Industrial Agriculture (MINEPIA) inspectors carried out a postmortem on the carcass, looking for evidence of granulomatous bTB-like lesions and evidence of hepatic fibrosis by *Fasciola gigantica* (liver fluke). If an animal had bTB-like lesions then up to 3 lesions per animal were collected for culture and a random sample of retropharyngeal lymph nodes from non-lesioned animals was also collected for comparison. More in depth detail about the study design and diagnostic tests carried out in this study can be found in Egbe et al. (2016, 2017) and Kelly et al. (2018). Samples with lesions from which *M. bovis* was recovered in addition to a randomly selection of controls from samples that were free of the disease, were archived (n=258, Egbe et al. (2016, 2017)). Selection took into account the age structure and breed structure in each of the abattoirs.

**Figure 1.**
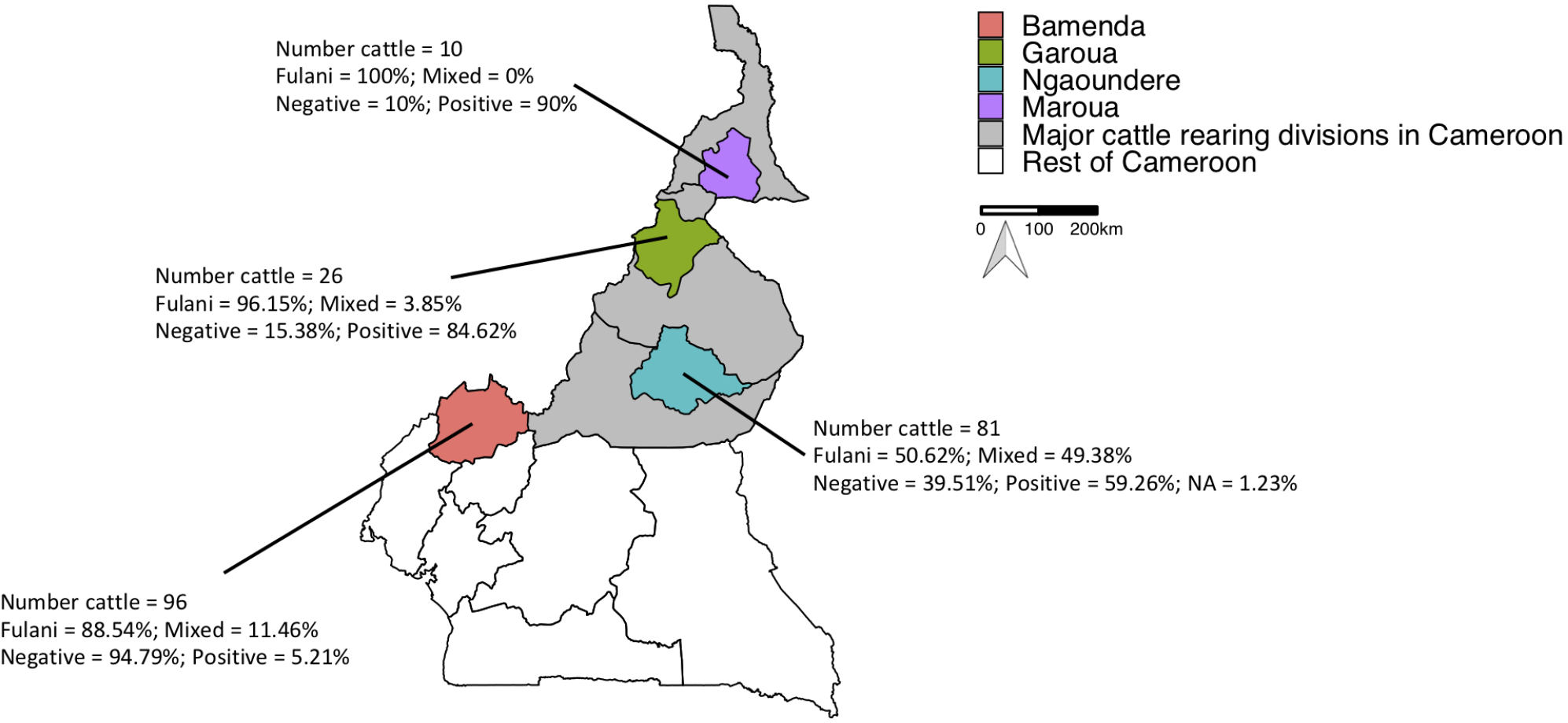
Map of Cameroon showing the major cattle rearing divisions and the divisions where the abattoirs are located. The number of cattle per abattoir, the proportion of mixed and Fulani cattle, and proportion of *M. bovis* infected cases per abattoir are also shown.

In this paper, the archived samples were genotyped. Cases were defined as *M. bovis* infection positive animals. Cases had at least one lesion which was culture positive using Mycobacterial Growth Indicator Tubes (MGIT), Lowenstein Jensen (LJ)-pyruvate or LJ-glycerol and typed positive for *M. bovis* (n=84 genotyped animals retained after quality control of the Illumina reads). Controls were animals which were lesion and culture negative for *M. bovis* (n=128 genotyped animals retained after quality control of the Illumina reads). An additional Fulani animal was also genotyped but we did not have any information on its *M. bovis* type infection status, although it was both lesion and culture positive.

### 2.2 Culture and bacteriological typing

Inoculum was prepared from lymph nodes (Egbe et al., 2016, 2017) and a portion cultured in Mycobacterial Growth Indicator Tubes (MGIT), these were incubated for 8 weeks on the BACTEC MGIT 960 automated culture system (Becton, Dickinson and Company, 1 Becton Drive, Franklin Lakes, NJ, USA) following the manufacturer’s instructions. Negative MGITs were examined visually after 56 days for growth before discarded (Egbe et al., 2017). Another portion of the inoculum was cultured on Lowenstein Jensen (LJ) media (one supplemented with pyruvate and the other with glycerol) and observed weekly for up to 12 weeks. After 12 weeks, if no growth was observed then they were classified as culture negative (Egbe et al., 2016). If any growth was suspected, on both media, a smear was prepared, stained according to the Ziehl-Neelsen (ZN) method and examined with a microscope (100x magnification) to assess the presence of acid-fast bacilli (Lumb et al., 2013). All samples with observed acid-fast bacilli were screened for the presence of *M. bovis* using the Hain GenoType^®^ MTBC assay (Hain Lifescience^®^, GmbH, Nehren, Germany) (Egbe et al., 2016, 2017). *M. bovis* cases are therefore defined as animals that have one or more lesions which were positive by one or more culture methods and were confirmed by typing using the Hain GenoType^®^ MTBC assay.

### 2.3 Genotyping and quality control

DNA was extracted from the frozen archived bovine lymph nodes using the Qiagen DNeasy^®^ Blood & Tissue Kit. The ‘Purification of Total DNA from Animal Tissues (Spin-Column Protocol)’ was used following the manufacturer’s instructions.

Frozen lymph node samples stored at −80°C were brought to room temperature and approximately 25mg was sliced into tiny pieces on a sterile petri dish using sterile disposable surgical blades. The sliced tissues were transferred to a 1.5ml screw cap microtube and heat-treated for 10min at 95°C in a water bath. Tubes were retrieved from the water bath and 180 μl of Buffer ATL and 20 μl of proteinase K (provided with the kit) was added. Tubes were vigorously vortexed to mix and the tissues were completely lysed by incubating at 56°C for 3 hours in a heater block. The rest of the procedure was done strictly following the manufacturer’s instructions. Purified DNA was first eluted from the spin columns with 60 μl of the elution buffer to increase the DNA concentration. To maximise the DNA yield, a second elution with 180 μl of elution buffer was done. The concentration of the purified DNA were assessed using the Qubit.

The cattle were genotyped using the Illumina BovineHD 777K BeadChip, which included 777,962 SNPs, of which 735,293 were autosomal (Illumina, 2015). In the archived subset there were lymph node tissue samples from 172 Fulani cattle and 56 mixed breeds. In order to assess the genomic architecture in *Bos indicus* breeds in Cameroon, sequences for an additional 216 reference animals genotyped using the Illumina BovineHD 777K BeadChip, were obtained. These reference animals represent European taurine breeds (Holstein n=63 and Jersey n=36), African taurine breeds (N’Dama n=24, Muturu=10), Asian zebu breeds (Nelore n=35, Gir=30) and an East African admixed breed (Sheko n=18). The details of the origins of these animals are provided in Table S1.

Quality control of the reads was carried out using the program PLINK v1.90 (Purcell and Chang, 2018; Chang et al., 2015). SNPs with a minor allele frequency (MAF) of <0.01 or a call rate of <90% were removed. Unless stated, no Hardy-Weinberg equilibrium cut-off was used to avoid removal of informative SNPs. Individuals that were related with a degree of relatedness greater than two according to the KING relatedness algorithm were removed (Manichaikul et al., 2010). This left a total of 500815 autosomal SNPs and 320 cattle, of which 213 animals were from Cameroon.

#### *Bos indicus* genetic diversity

Investigation of the *Bos indicus* genetic diversity was carried out by comparing the Fulani and the mixed breed animals from Cameroon (n=213) to 107 reference animals that passed quality control checks (Table S1).

#### Population genetic structure

Principle components analysis (PCA) was performed using the *pca* function of PLINK v1.90 (Purcell and Chang, 2018; Chang et al., 2015) to provide an insight into the population structure of the cattle breeds.

Next, hierarchical clustering was performed on the genome-wide identity-by-state (IBS) pairwise distances between individuals using the *SNPRelate* package in R version 3.5.0 (Zheng et al., 2012; R Core Team, 2018). Subgroups of individuals were determined using a Z-score threshold of 15 based upon individual dissimilarities to define groups of individuals in the hierarchical cluster analysis. An outlier threshold of 5 was also set, this means that groups with less than or equal to 5 animals are considered outliers. For comparison the dendrogram was redrawn using breed and population type to determine the groups.

Population genetic structure was also evaluated using the ADMIXTURE software tool (Alexander et al., 2009) to determine the European taurine, Asian zebu and African taurine ancestries at the genome-wide level. Variants in high linkage disequilibrium (LD) with each other were removed prior to analysis. The LD pruning criteria applied was to remove any SNP that had an r-squared >0.2 with another SNP within a 200-SNP window; for a sliding window of 10 SNPs at a time, which resulted in 55132 markers (out of 500815 markers) left for analysis. A 5-step expectation–maximisation (EM) algorithm was used. In addition, 10-fold cross validation was performed with 200 bootstrap resampling runs to estimate the standard errors for each cluster level (K=2 to 12). The output was plotted using the *pophelper* package for R (Francis, 2017). The optimal number of clusters was determined from the cross-validation plot.

#### Genetic differentiation and inbreeding coefficients

To estimate the degree of genetic differentiation in the cattle populations, fixation indices (Fst) were calculated using the Weir and Hill (2002) relative beta estimator method as implemented by the *snpgdsFst* function in the *SNPRelate* package (Zheng et al., 2012; Weir and Hill, 2002; Buckleton et al., 2016). In addition, the *het* function of PLINK (Purcell and Chang, 2018) was used to calculate the inbreeding coefficient estimate, F, for each individual.

### Genetics of bTB resistance

#### Association between bTB infection status and breed

Generalised linear models in R (R Core Team, 2018) were used to examine the association between breed and bTB infection status. Two separate models were run, the first used the local abattoir employees record of breed to test the association between breed and bTB. The model structure for this analysis was the same as that used in Kelly et al. (2018) which accounted for age and sex as fixed effects, however, this time the analysis was based on the 212 genotyped cattle instead of the 935 phenotyped cattle (Kelly et al., 2018). The second method used the same model structure but breed was replaced with the hierarchical clustering definition of subgroups of animals based on genomic information. For all models, logit link functions were used. It was not possible to include abattoir as additional fixed effects in these models as some abattoirs did not contribute enough individuals with complete data. It is noted that, excluding these potential confounders could have large effects on the estimates, as they may absorb all other potential systematic environmental effects on bTB status.

#### Association with inbreeding and European taurine introgression

European taurine (ET) introgression status was based on the proportion membership to the best supported K value cluster representing European Taurine breeds in the admixture analysis (Cluster 3 from K=6). We defined ‘moderate’ ET introgressed cattle as individuals with between 1% and 10% ET introgression (n=72) and cattle with ≤1% ET background represented the ‘non-European’ introgressed group (n=141). There were no ‘substantially’ ET introgressed cattle with >10% ET introgression in this population.

Cattle were defined as inbred using a method adapted from Murray et al. (2013), which we described here. Non-ET introgressed cattle were categorised as inbred if they had an inbreeding coefficient value of greater than 0.184 (more than 0.10 above the mean for this group). As in Murray et al. (2013), for ET introgressed animals, the best fit linear regression of inbreeding against introgression was found and cattle were excluded if they had an inbreeding coefficient >0.01 above their expected value. This resulted in the identification of 21 inbred cattle, 17 of which showed moderate ET introgression.

We examined the relationship between bTB and (i) ET introgression and (ii) the inbreeding coefficient using generalised linear models with logit link functions in R (R Core Team, 2018). A separate model was built for each. Age and sex were included in the models as categorical fixed effects. Again, it was not possible to include abattoir in these models as some abattoirs did not have any inbred or ET introgressed cattle.

#### Heritability analysis

The heritability of bTB resistance in the Cameroon dataset was assessed using linear animal models (Lynch et al., 1998), which are a form of mixed model with fixed and random effects, that can break phenotypic variation down into the different components using the following structure:

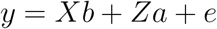

where *y* is the phenotype of interest (in this case, *M. bovis* infection status) and *b* is a vector of fixed effects (age, sex and breed). The random effects, which determine the variance of the trait are the additive genetic (*a*) and residual effects (*e*). *X* and *Z* are all design matrices assigning individuals to their corresponding fixed and random effects. Both random effects were assumed to follow a normal distributions, a 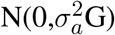, where 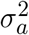 is the additive genetic variance and G is the genomic relationship matrix constructed from the inverse of the IBS matrix which was created using the R package *SNPRelate* (Zheng et al., 2012) as described above; the residual error e 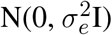 with residual variance 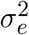 and identity matrix I.

The narrow-sense heritability of a trait (*h*^2^) is defined as the proportion of phenotypic variance explained by the additive genetic variance, 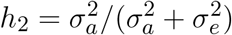. It describes the extent to which differences between individuals are determined by additive genetic effects (Falconer et al., 1996). The heritability analyses were carried out in ASReml version 3.0 (Gilmour et al., 2009). Note, these estimates are likely to be inflated as they do not include any population structure or other systematic environmental effects that may be confounded with genetic effects. It was not possible to include these in the model due to the small numbers of individuals genotyped from each abattoir in this study.

#### Genome wide association study

A genome wide association study (GWAS) was carried out on the quality controlled Cameroon dataset using PLINK version 1.90 (Purcell and Chang, 2018; Chang et al., 2015). Linear regression models were run to evaluate the association between bTB and each SNP. Age, sex and breed were included in the models as fixed effects. A Bonferroni correction was applied to the genome-wide significance threshold, calculated as *p* = −*log*_10_(0.05/*n*) = 7.00 (where *n* is the number of SNPs, *n* = 500815). Regions surrounding 100000bp of SNPs which had been identified as being associated with bTB resistance in cattle in the literature (Bermingham et al., 2014; Finlay et al., 2012; Richardson et al., 2016; le Roex et al., 2013; Tsairidou et al., 2018; Amos et al., 2013; Driscoll et al., 2011; Raphaka et al., 2017) were also evaluated with a lower significance threshold of *p* = −*log*_10_(0.05/45780) = 5.96.

To investigate breed differences in *M. bovis* infection status further, the minor allele frequency of SNPs associated with *M. bovis* infection status with a p value − 1 × 10^−5^ to account for multiple comparisons were compared between breeds.

## RESULTS

### Descriptive results

A total of 213 animals from Cameroon passed quality control checks, of which 161 animals (75.6%) were Fulani and 52 animals (24.4%) were mixed breed. 84 animals (39.4%) were positive for *M. bovis* infection (Table 1). Within the Fulani group 38.5% of animals (62 individuals) were positive for *M. bovis* infection compared to 42.3% of mixed breed animals (22 individuals). One Fulani animal was missing a *M. bovis* typing test result. The proportion of individuals positive for *M. bovis* infection by age, gender and abattoir is shown in Table 1.

**Table 1.** The number and percentage of cattle by breed and abattoir with *M. bovis* infection.

| Breed | Abattoir | Total |  | Female |  | Male |  | Young |  | Old |  | Inbred |  | Outbred |  | No European Taurine Introgression |  | Moderate European Taurine Introgression |  | *M. bovis* infection positive |  |
| --- | --- | --- | --- | --- | --- | --- | --- | --- | --- | --- | --- | --- | --- | --- | --- | --- | --- | --- | --- | --- | --- |
|  |  | N | % | N | % | N | % | N | % | N | % | N | % | N | % | N | % | N | % | N | % |
| Fulani | Bamenda | 85 | 89 | 35 | 41 | 50 | 59 | 29 | 34 | 56 | 66 | 14 | 16 | 71 | 84 | 52 | 61 | 33 | 39 | 5 | 6 |
| Fulani | Garoua | 25 | 96 | 20 | 80 | 2 | 8 | 0 | 0 | 25 | 100 | 0 | 0 | 25 | 100 | 22 | 88 | 3 | 12 | 21 | 84 |
| Fulani | Maroua | 10 | 100 | 9 | 90 | 1 | 10 | 0 | 0 | 10 | 100 | 0 | 0 | 10 | 100 | 6 | 60 | 4 | 40 | 9 | 90 |
| Fulani | Ngaoundere | 41 | 51 | 37 | 90 | 4 | 10 | 5 | 12 | 35 | 85 | 3 | 7 | 38 | 93 | 29 | 71 | 12 | 29 | 27 | 66 |
| Mixed | Bamenda | 11 | 11 | 1 | 9 | 10 | 91 | 2 | 18 | 9 | 82 | 1 | 9 | 10 | 91 | 5 | 45 | 6 | 55 | 0 | 0 |
| Mixed | Garoua | 1 | 4 | 0 | 0 | 0 | 0 | 0 | 0 | 1 | 100 | 0 | 0 | 1 | 100 | 0 | 0 | 1 | 100 | 1 | 100 |
| Mixed | Maroua | 0 | 0 | 0 | 0 | 0 | 0 | 0 | 0 | 0 | 0 | 0 | 0 | 0 | 0 | 0 | 0 | 0 | 0 | 0 | 0 |
| Mixed | Ngaoundere | 40 | 49 | 38 | 95 | 2 | 5 | 6 | 15 | 34 | 85 | 3 | 8 | 37 | 92 | 27 | 68 | 13 | 32 | 21 | 52 |
| Total |  | 213 | 100 | 140 | 65.7 | 69 | 32.4 | 42 | 19.7 | 170 | 79.8 | 21 | 10 | 192 | 90 | 141 | 66 | 72 | 34 | 84 | 39.4 |

### *Bos indicus* genetic diversity

#### Population structure

PCA and hierarchical clustering analysis were used to show the population structure of the cattle breeds. Within the Cameroon cattle population, the PCA showed that the Fulani and mixed breed cattle cluster closely together along with the admixed Sheko cattle (Figure 2). The African taurine, European taurine breeds and Asian zebu breed form their own distinct clusters. The first two components of the PCA account for 53.1% and 15.1% of the total variation, respectively.

**Figure 2.**
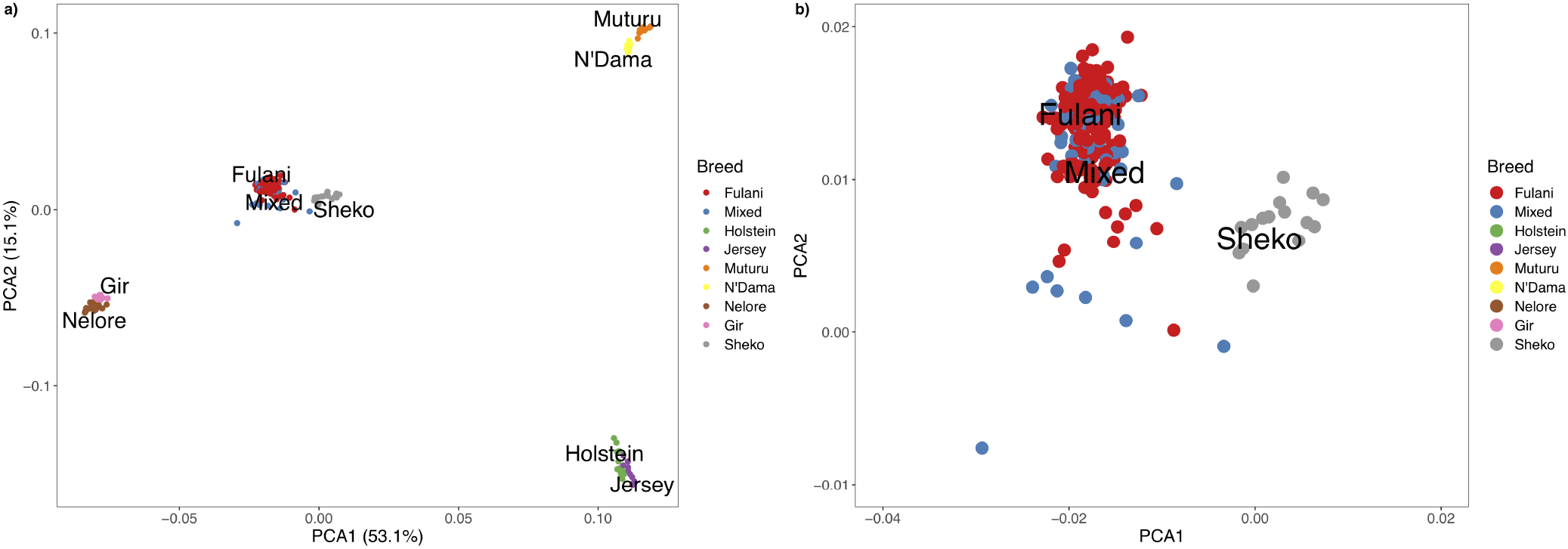
a) PCA plot for the Fulani and mixed breeds with the reference cattle breeds for the first two principle components. b) Zoomed in PCA plot showing the Fulani and mixed breeds.

The hierarchical clustering identified 10 large clusters of animals, shown by the grey and white bands and similarly coloured nodes in Figure 3a. The remaining clusters were outliers, consisting of less then 5 animals in each group, and are highlighted by the red bands in Figure 3a. In Figure 3b, the same hierarchical relationship amongst individuals is shown as in Figure 3a, apart from the nodes are shaded according to the local abattoir employees definition of breed rather than cluster membership. In combination, Figure 3a and Figure 3b confirm the PCA that the mixed breed cattle are grouped in the same cluster as the Fulani cattle and the admixed Sheko.

**Figure 3.**
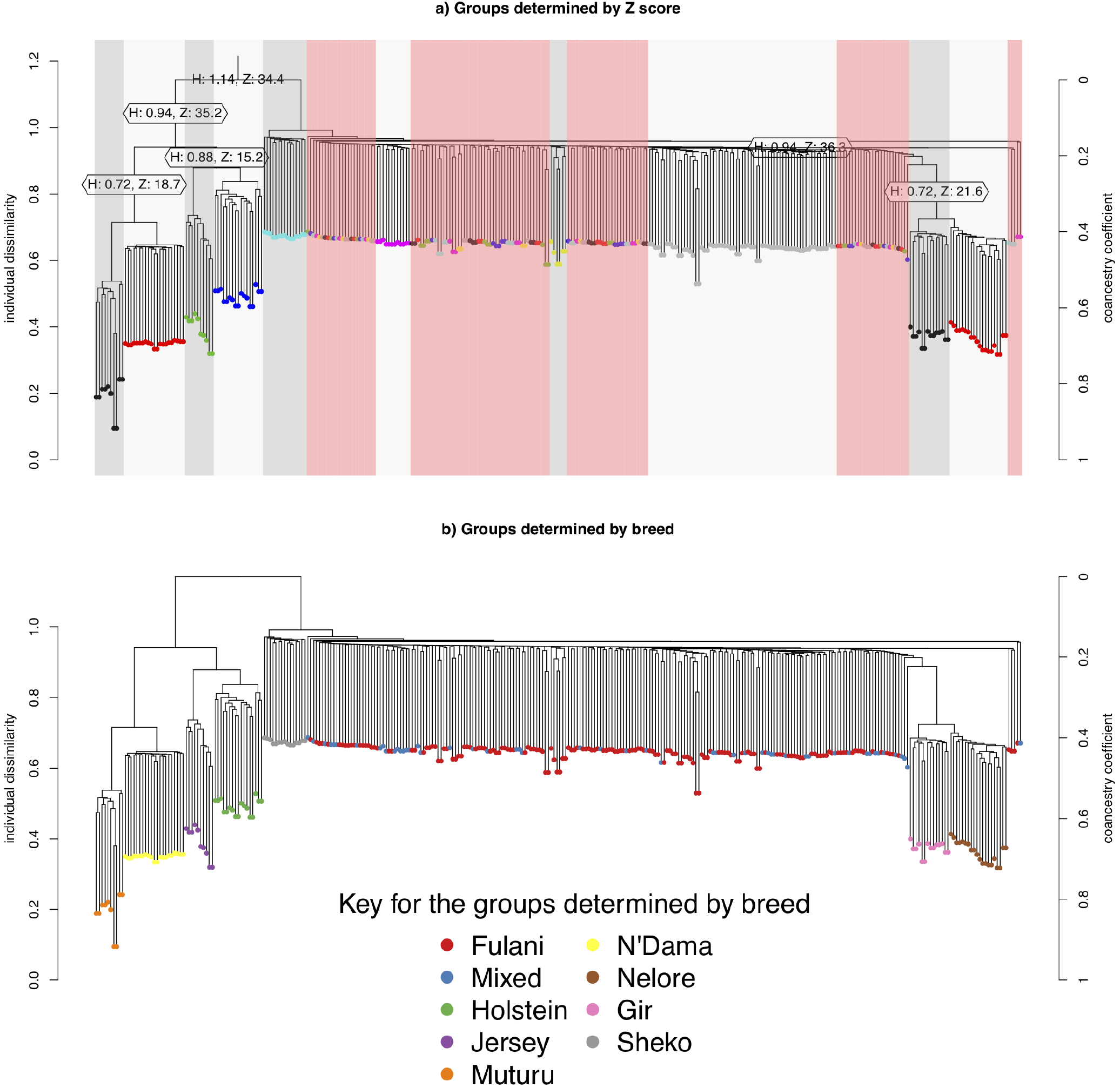
Hierarchical clustering of the IBS matrix. (a) Shows the groups as determined by individual dissimilarity and z-score, the colour of the nodes represent membership to each of the clusters using a z-threshold of 15. The grey and white highlighted areas symbolise the cluster groups, each labelled with their unique cluster name. The red highlighted areas represent the outliers, which are made up of clusters with less than 5 individuals in them. (b) Shows the hierarchical relationship amongst individuals, the nodes are shaded according to the local abattoir employee’s definition of breed

The population structure was investigated further using ADMIXTURE, with K=2 to K=6 shown in Figure 4. At K=2 the cattle are split into the indicus and taurine groups. At K=3, the African taurine and European taurine cattle diverge. In K=4 to K=6, the admixed cattle diverge, and higher levels of genetic heterogeneity in the admixed breeds compared to the indicus and taurine breeds is observed (Figure 4). Inspection of the cross-validation plot (Figure S1) suggests that K=6 is the optimal cluster number to describe the ancestry in this population. At K=6, the Fulani and mixed cattle show similar levels of heterogeneity with some cattle having a larger proportion of indicus admixture then others (Figure 4). There is very little European taurine introgression in either the Fulani or mixed cattle.

**Figure 4.**
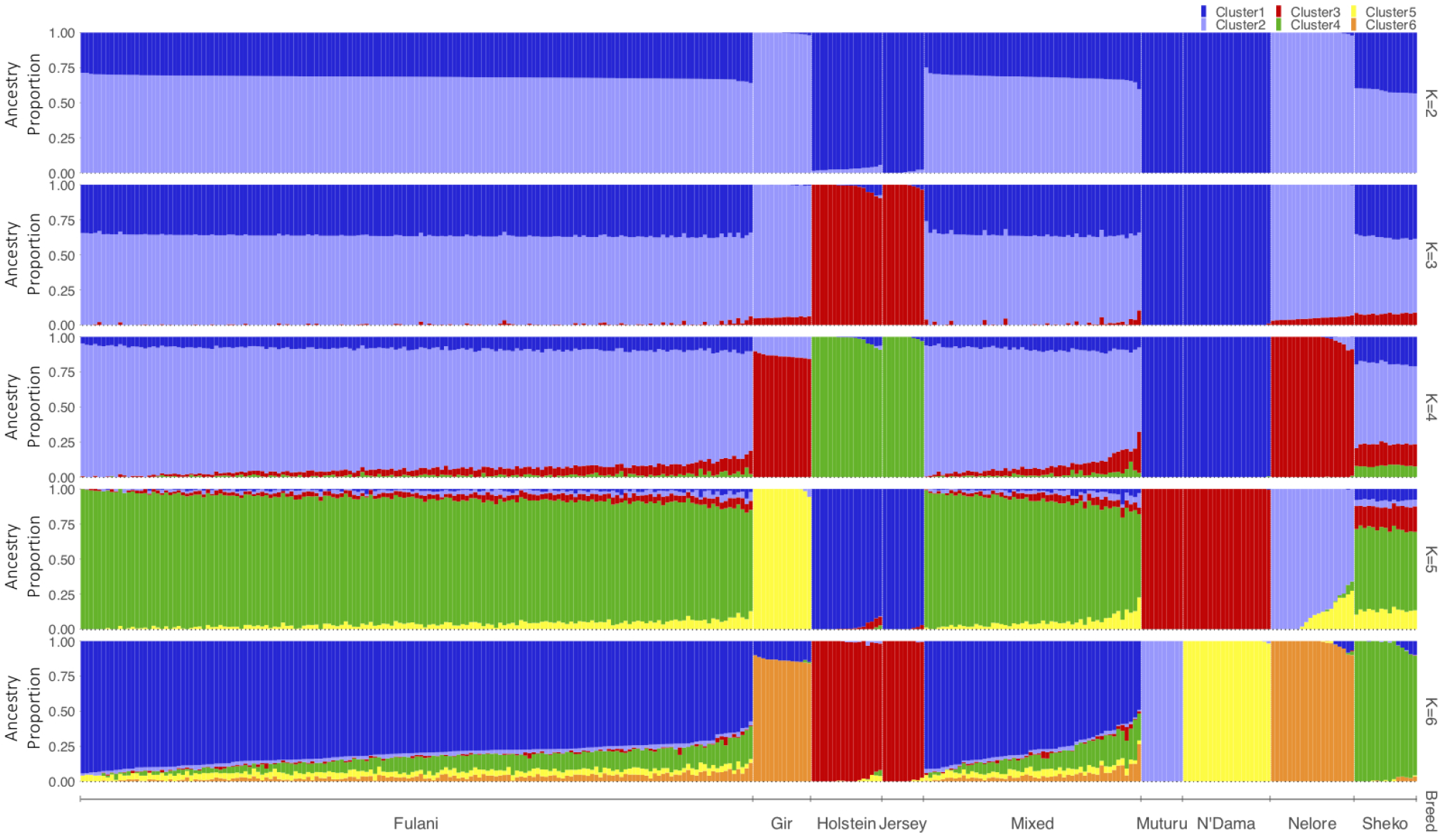
Admixture bar plots for the proportion of genetic membership to each ancestry assuming (K=2 to K=6) ancestral populations. Each animal is represented by a vertical line divided into K colours, indicating the likelihood of the animals genome belonging to an ancestral population

#### Genetic differentiation and inbreeding coefficients

High levels of genetic differentiation were observed between all the different breeds (Fst=0.284; Mean Fst=0.271; Range Fst = −0.071 - 0.991), however focusing on just the mixed and Fulani breeds there were low levels of genetic differentiation (Fst=0.001; Mean Fst=0.001; Range Fst = −0.009 - 0.14) indicating that there are high levels of shared genetic material between the Fulani and mixed breed animals (Figure S2).

The observed inbreeding coefficient, F is shown in Figure 5. Out of the animals we studied, the Muturu animals (which are an African taurine breed) have the highest level of inbreeding (mean F= 0.563, SE=0.011) whilst the admixed Sheko animals were the most outbred group (mean F= 0.037, SE=0.003). The Fulani cattle had a mean inbreeding coefficient of 0.082 (SE= 0.003) in comparison to 0.072 for the mixed cattle (SE=0.004).

**Figure 5.**
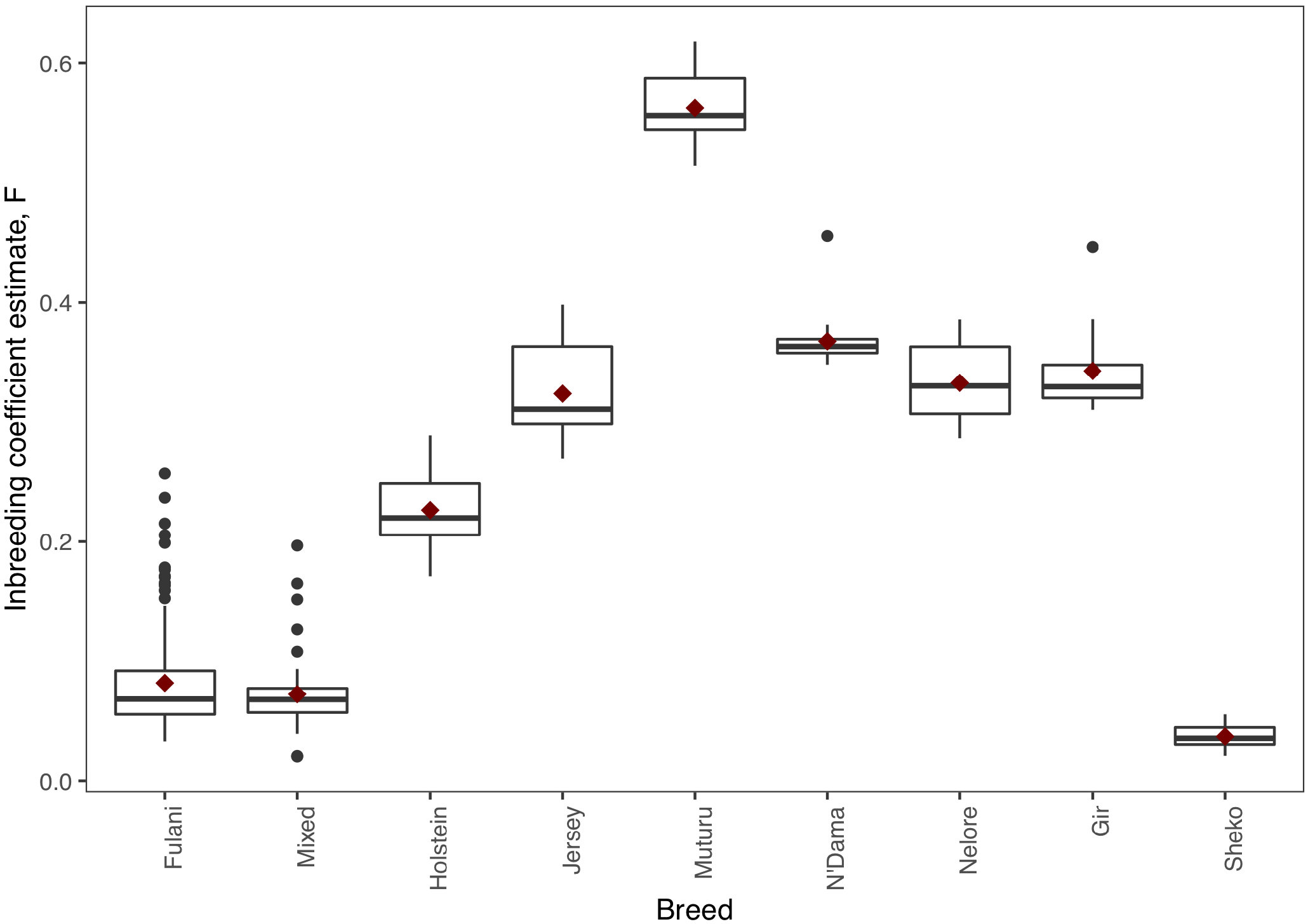
The observed inbreeding coefficient estimate, F, for each animal by breed. The red diamond represents the mean value for each breed.

### bTB resistance genetics

Kelly et al. (2018) observed that there was an association between bTB lesions and breed as defined by the local abattoir employees. In our model which looked at the association between *M. bovis* infection status and the local abattoir employees definition of breed, only accounting for age and sex, there was no association with breed in the genotyped population (Table 2). Furthermore, when we model the association between *M. bovis* infection and breed using the same model structure as in Kelly et al. (2018), breed is only associated with *M. bovis* infection status when *Fasciola* sp. infection status is included in the model (Table S2).

**Table 2.**
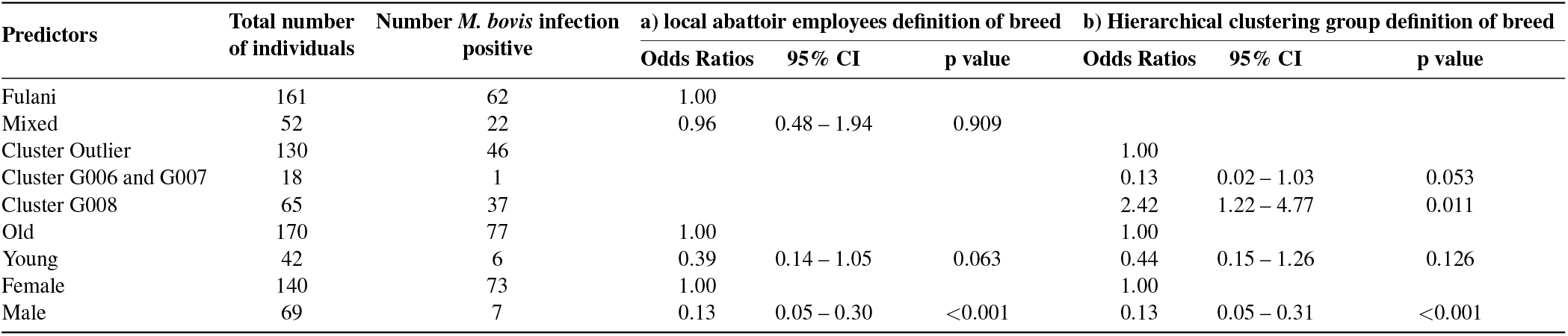
Association between *M. bovis* infection and (a) local abattoir employees definition of breed and (b) hierarchical clustering group definition of breed after accounting for age and sex (n=207)

However, when membership to hierarchical clustering group is used instead of the local abattoir employee’s definition of breed, there was a difference in risk of *M. bovis* infection between groups, Figure 6. Cattle belonging to cluster G008 were more than twice as likely of being *M. bovis* infection positive compared to cattle in the ‘outlier’ cluster after accounting for age and sex (OR=2.42, 95% CI = 1.22-4.77, Table 2) whilst there was no difference in risk for cattle in cluster G006/G007 and the outlier cattle (OR=0.13, 95% CI = 0.02-1.03, Table 2).

**Figure 6.**
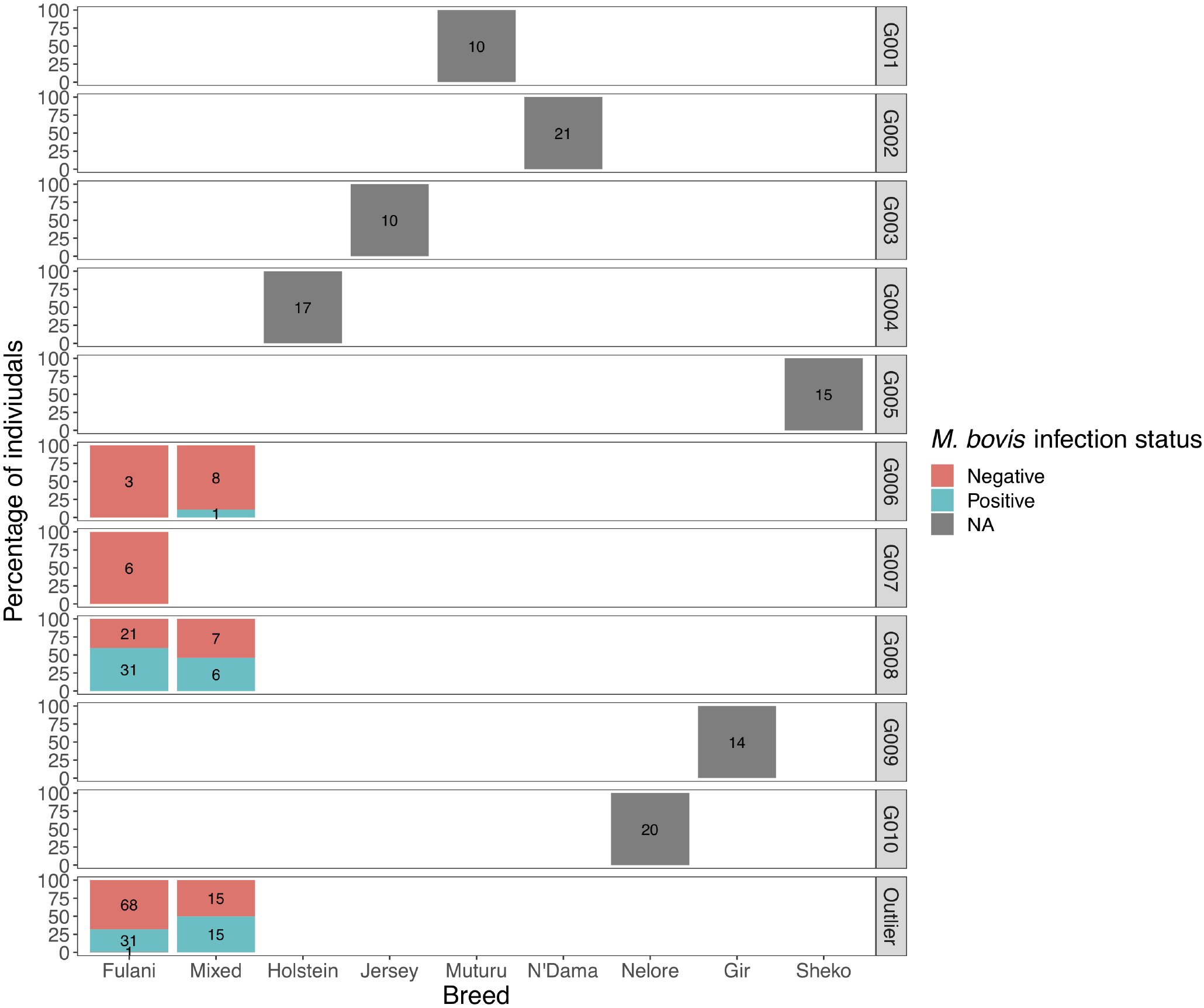
The percentage of individuals belonging to each hierarchical cluster group as shown in Figure 3 broken down by the local abattoir employees definition of breed and *M. bovis* infection status. The labels refer to number of individuals belonging to each group

The characteristics of each cluster for the Fulani and mixed animals are described in Fig. 7. There was no difference between cattle in G008 and the outliers in terms of the local abattoir employees definition of breed, abattoir or inbreeding status. However, cattle in the outlier cluster were more likely to be ET introgressed then the G008 cattle (OR = 3.14, 95% CI = 1.57 - 6.67, p=0.002 for the risk of moderate ET introgression in the outlier cattle in comparison to G008 animals).

**Figure 7.**
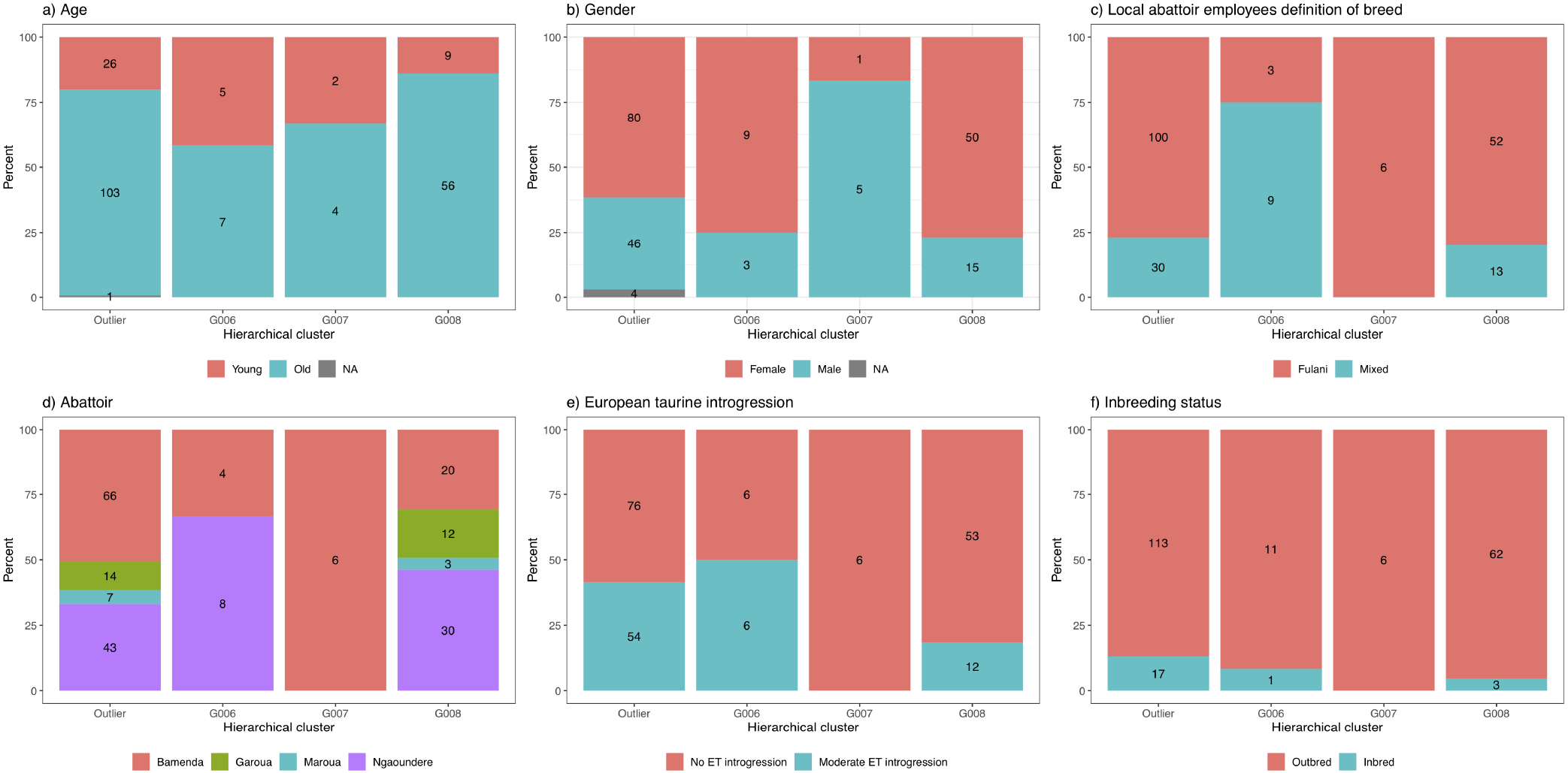
The number and percentage of animals in each hierarchical cluster according to (a) age, (b) gender, (c) local abattoir employees definition of breed, (d) abattoir, (e) European taurine introgression and (f) inbreeding status. The labels refer to number of individuals belong to each group

Yet, in separate generalised linear models after accounting for age and sex there was no association between European taurine introgression and *M. bovis* infection status (Table 3). In contrast, a further generalised linear model showed that inbred cattle have 0.15 times the odds of having *M. bovis* infection than outbred cattle, this means that inbred cattle were significantly less likely to be *M.bovis* positive than outbred cattle (OR=0.15, 95% CI= 0.03-0.68, p=0.014, Table 3).

**Table 3.**
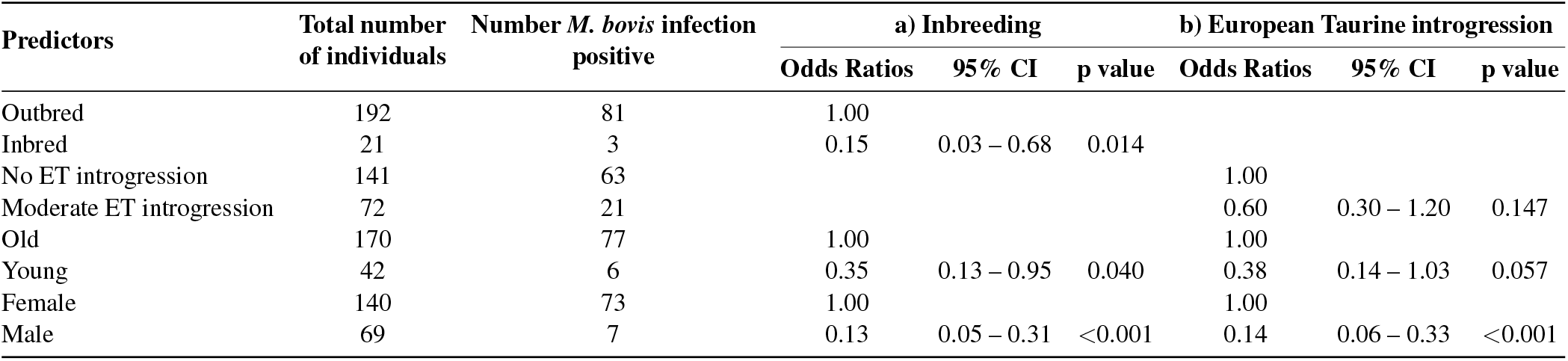
Association between (a) inbreeding and (b) European Taurine introgression and *M. bovis* infection status after accounting for age and sex (n=207)

| Predictors | Total number of individuals | Number *M. bovis* infection positive | a) Inbreeding |  |  | b) European Taurine introgression |  |  |
| --- | --- | --- | --- | --- | --- | --- | --- | --- |
|  |  |  | Odds Ratios | 95% CI | p value | Odds Ratios | 95% CI | p value |
| Outbred | 192 | 81 | 1.00 |  |  |  |  |  |
| Inbred | 21 | 3 | 0.15 | 0.03 – 0.68 | 0.014 |  |  |  |
| No ET introgression | 141 | 63 |  |  |  | 1.00 |  |  |
| Moderate ET introgression | 72 | 21 |  |  |  | 0.60 | 0.30 – 1.20 | 0.147 |
| Old | 170 | 77 | 1.00 |  |  | 1.00 |  |  |
| Young | 42 | 6 | 0.35 | 0.13 – 0.95 | 0.040 | 0.38 | 0.14 – 1.03 | 0.057 |
| Female | 140 | 73 | 1.00 |  |  | 1.00 |  |  |
| Male | 69 | 7 | 0.13 | 0.05 – 0.31 | <0.001 | 0.14 | 0.06 – 0.33 | <0.001 |

#### Heritability analysis and genome wide association study

The crude heritability of being *M. bovis* infection positive in the Cameroon population, after accounting for age, sex and breed is *h*^2^=19.8% (SE=16.5, Table S3). After accounting for age, sex and breed, there is a suggestion of an association between SNPs on Chromosome 12 (region 34703645 - 34780062) and *M. bovis* infection status at the suggestive threshold rather than the genome wide threshold (P=< 1 × 10^−5^, Figure 8 and Figure S3 for the QQ-plot). None of these SNPs were in regions which had been previously identified as associated with bTB resistance in cattle in the literature. Ensembl release 97 (Zerbino et al., 2017) identified five genes within ±500Kbp of this region (12:34703645 - 34780062): mitochondrial intermediate peptidase (MIPEP); sacsin molecular chaperone (SACS); spermatogenesis associated 13 (SPATA13); TNF receptor superfamily member 19 and C1q and TNF related 9 (C1QTNF9).

**Figure 8.**
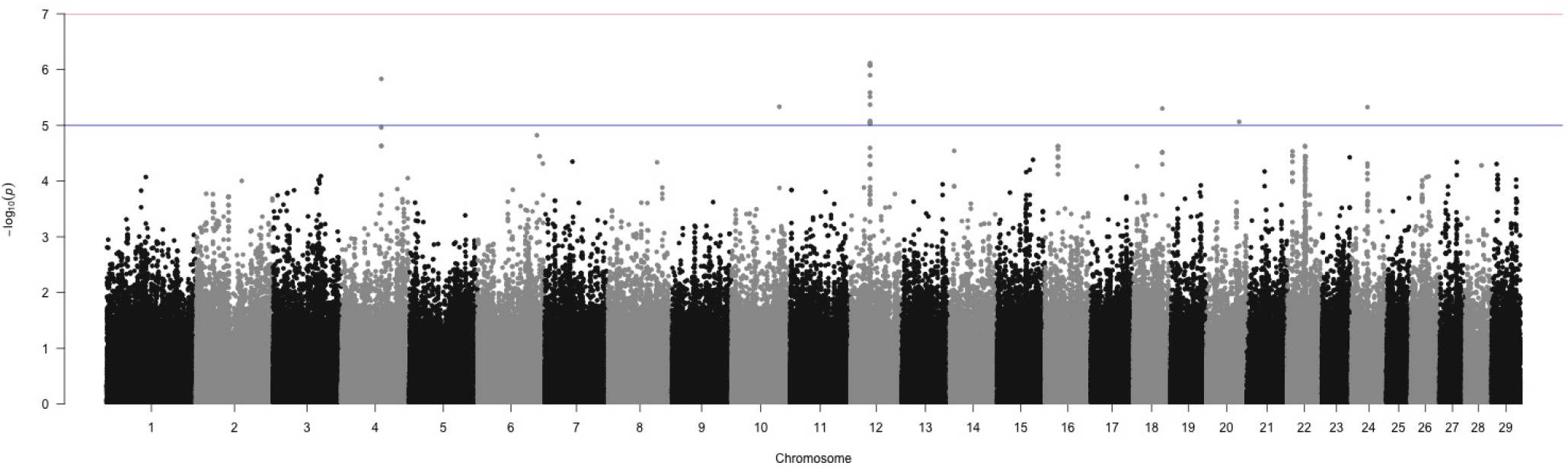
Manhattan plot of the genome wide association between SNPs and *M. bovis* infection status after accounting for age, sex and breed as covariates. The blue line represents the suggestive significance line of < 1 × 10’^5^ and the red line represents the genome wide significance threshold, using a Bonferroni correction of −*log*_10_(0.05/*n*).

The minor allele frequency (MAF) of the SNPs associated with *M. bovis* infection after accounting for the covariates age, breed and sex with a p value < 1 × 10^−5^ (n=55) cluster-stratified by breed is shown in Figure 9. There is no difference in the MAF of these SNPs in Fulani and mixed cattle, however the median MAF of these SNPs is lower in Asian zebu cattle compared to European and African taurine cattle (Figure 9). The admixed cattle have intermediate MAF for these SNPs.

**Figure 9.**
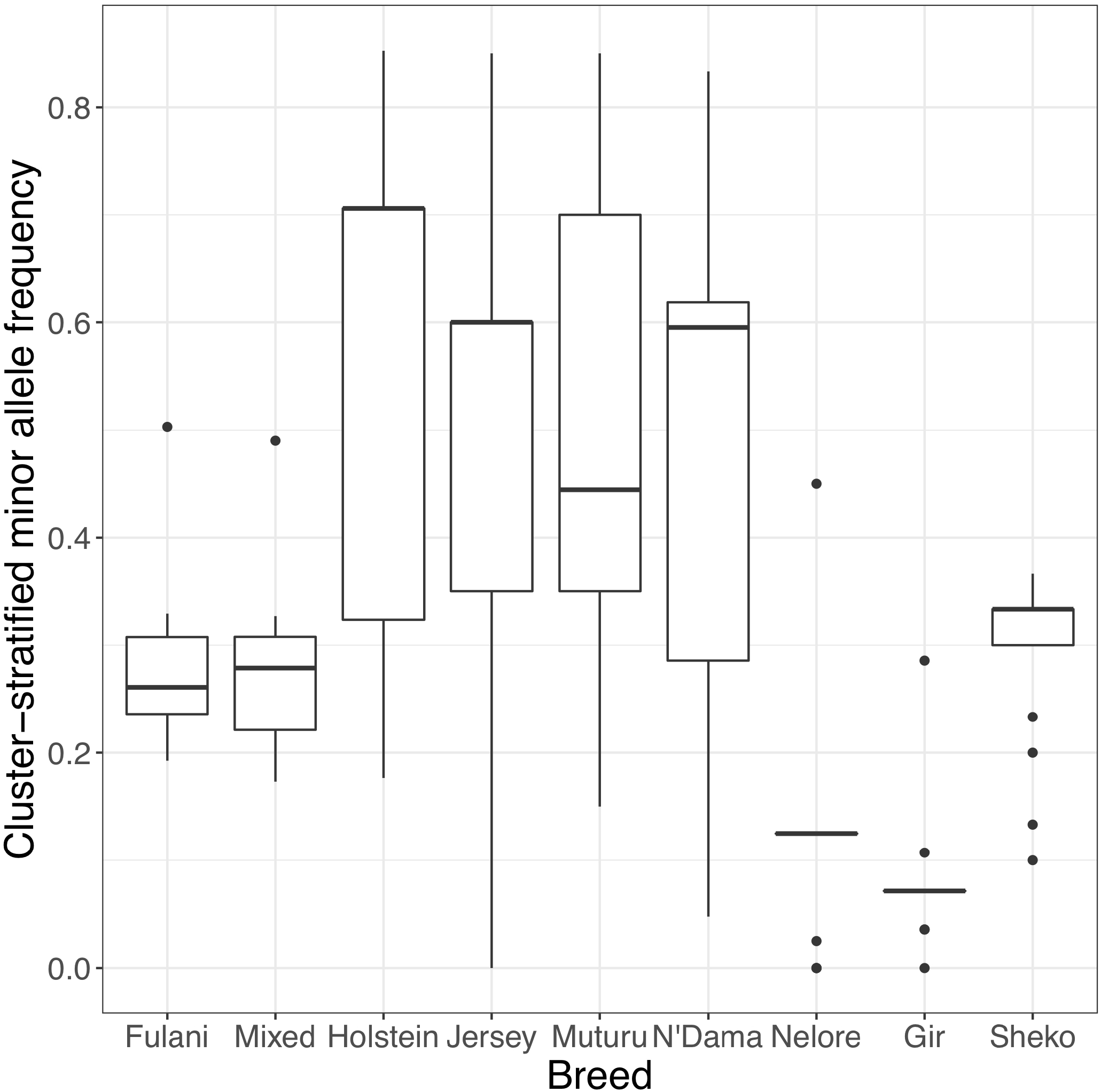
Cluster-stratified minor allele frequencies of the SNPs associated with *M. bovis* infection status with a p value < 1 × 10^−5^ by breed

## DISCUSSION

This study was aimed at reconciling the difference between the phenotype and genotype of the Fulani and mixed cattle in Cameroon following on from our previous study (Kelly et al., 2018). Although the local abattoir employees could identify a difference between the breeds, the apparent phenotypic differences in visual appearance between the breeds were not reflected by clear genomic differences. Within the Cameroon cattle population, the PCA showed that the Fulani and mixed breed cattle cluster closely together and the hierarchical clustering also grouped them together. Furthermore, the Fst analysis quantified the level of genetic differentiation between the two as very low (Fst=0.001, Mean Fst=0.001; Range Fst = −0.009 - 0.14) implying that the Fulani and mixed cattle are not genetically distinct. They also had similar levels of heterogeneity in the admixture analysis.

A study in East Africa has also compared farmer and field staff assessments of breed, based on phenotypic appearance with the admixture determinations of breed composition. They showed that that phenotype-based assessments were very poor predictors of actual breed composition (*R*^2^ = 0.16) (Marshall et al., 2019).

Reasons for this discrepancy could be because breed may be a proxy for husbandry. Breed names can relate to the location where animals are found or ethnic group they are kept by rather than genetic differences between breeds (Rege and Tawah, 1999; Frantz, 1993; Burnham, 1996). Husbandry factors such as pooled management (keeping all animals together and using bulls of the same breed) have also been attributed to the low levels of genetic differentiation found between Fulani cattle (Ibeagha-Awemu and Erhardt, 2006).

Alternatively, the marker diversity of the SNP chip may not accurately reflect the genetic diversity of the *Bos indicus* breeds. The Illumina BovineHD 777K BeadChip was validated in economically important European beef and dairy cattle (Illumina, 2015) and may lack the discriminatory power to differentiate between different *Bos indicus* breeds. It may be missing variants responsible for disease resistance as well as observed physical characteristics. In the future, this analysis should be repeated with a custom SNP chip designed specifically for *Bos indicus* cattle to rule this out. These are currently being developed by the Centre of Tropical Livestock Genetics and Health (CTLGH) who are currently genotyping hundreds of cattle to generate a new SNP chip for the Africa content which prioritises SNPs for key traits such as bTB resistance.

There was very little European taurine introgression in both the Fulani and mixed cattle. Unlike other African countries, there has been very limited introduction of European breeds into Cameroon (Muwonge et al., 2019). Fulani cattle are kept by pastoral communities for dual purpose (meat and milk) and this lack of differentiation means that such a breed was less likely to be targeted for breed improvements by cross breeding with European taurine breeds. In addition, transhumance, the seasonal movement of livestock between pastures during the dry and wet season, practised by pastoralists in this region is dominated by native zebu breeds (Motta et al., 2018). Transhumance requires hardy, resilient and disease tolerant animals that can trek hundreds of miles along the Sahel transhumance highway (Motta et al., 2018; Turner and Schlecht, 2019). This limits the chances of breeding outside this gene pool since the characteristics targeted by breed improvements in most African settings are likely incompatible with the transhumance way of life.

Recently there have been introductions of exotic breeds and other taurine breeds into dairy improvement programs in Cameroon (Tambi, 1991; Njwe et al., 2002). These recent introductions will not be picked up in our sample of abattoir cattle as dairy cattle in Cameroon are very restricted in numbers and managed by small holders, who infrequently slaughter cattle or trade cattle with pastoralist groups (Kelly et al., 2016).

The literature is riddled with reports calling for the introduction of exotic breeds to be carefully managed. If uncontrolled cross-breeding continues without fully examining the socio-ecological, economical and environmental impact there is a risk of eroding the unique genetic resource which is adapted to the sub-Saharan environment (Mwai et al., 2015; Ibeagha-Awemu and Erhardt, 2006). The Fulani breeds are not only adapted for the transhumance way of life and the harsh climate (Hansen, 2004), it is known that African *Bos taurus* have unique evolutionary adaptations to endemic diseases. For example, they are tolerant to trypanosomosis and *Theileria parva* which causes East Coast fever (Roberts and Gray, 1973; Coetzer and Tustin, 2004).

We showed that cattle in hierarchical clustering group G008 were more likely to be *M. bovis* infection positive than cattle in the ‘outlier’ cluster. There was no difference in prevalence of *M. bovis* infection between individuals from G006 and G007 combined and the ‘outlier’ cattle. Individuals in cluster G008 could be found across the whole study area, they were not restricted to one abattoir and they were equally likely to be Fulani or mixed breed. Cattle in the outlier cluster were however more likely to be moderately ET introgressed then the ‘outlier’ cattle. Although preliminary and based upon small sample sizes, this result suggests that this attribute is ubiquitous in Cameroonian Fulani breeds and ought to be avoided to improve bTB control in such settings.

Furthermore, we found that inbred cattle were less likely to have *M. bovis* infection than outbred cattle. The reasons behind this are not so clear. It is possible that the small numbers (there were only 3 *M. bovis* infection positive inbred animals) could be driving this relationship. However, with more data it may be worth investigating further. Many studies, including studies of cattle, show the opposite effect, that inbreeding depression has negative effects on host fitness (Coltman et al., 1999; Leroy, 2014; Murray et al., 2013). Alternatively, it is possible that the inbred cattle, are less likely to be cross-bred with exotic breeds so therefore they are less likely to become infected with bTB.

Kelly et al. (2018) showed that there was increased risk of having bTB-like lesions in Fulani cattle compared to the mixed breed group using the local abattoir employee’s definition of breed in their study. Our paper only found an association between breed and bTB when including *Fasciola* sp. infection status in the model. There were a number of differences between the two analyses; the first was we only used a subset of individuals from the original Kelly *et al*. 2018 study. Secondly, we used individuals which where confirmed to have *M. bovis* infection by the Hain GenoType^®^ rather then presence of bTB lesions to determine bTB status, so our case definition is more specific. Alternatively, it could be the presence of *Fasciola* sp. which is driving the relationship. To unpick this relationship, all the cattle would need to be genotyped which is beyond the logistical scope of this current study but merits further investigation when resources allow.

We found no association between European taurine introgression and *M. bovis* infection status. However, Murray et al. (2013) found that Kenyan cattle with higher levels of European taurine introgression experienced more clinical illness. The difference observed here could be due to the substantially lower levels of introgression observed in Cameroon cattle (72 animals with 1-10% ET introgression), compared to the 113 animals with 12.5-36.1% ET introgression in Murray et al. (2013). Yet, the literature suggests that African zebu cattle are more resistant to bTB than exotic breeds of cattle (Vordermeier et al., 2012; Ameni et al., 2007).

In a cross-sectional study of bTB in Ethiopia, exotic cattle were more likely to be bTB positive using the comparative cervical intradermal tuberculin test (CIDT) than local cattle breeds (Habitu et al., 2019). A meta-analysis of bTB in Ethiopia also found that Holstein-Frisians have a higher bTB prevalence than local zebu (prevalence = 21.6% (95% CI: 14.7–30.7); prevalence = 4.1% (95% CI: 3.4–4.9), respectively) (Sibhat et al., 2017). Furthermore, Ameni *et al*. (Ameni et al., 2007) found that Holsteins had more severe bTB pathology than zebu cattle in Ethiopia. Therefore the use of and/or crossbreeding with taurine dairy cattle and the intensification of farming is likely to increase the incidence of bTB (Habitu et al., 2019).

Lastly, we investigated the role of additive genetic effects and individual SNPs on bTB resistance. We found that the heritability of bTB after accounting for age, sex and breed in this study was *h*^2^ = 19.8% (SE=16.5). The small sample size used for this heritability calculation has resulted in the large standard error.

It is possible that bTB resistance is polygenic with small penetrance at each gene. This means that there are few loci of large effect which partially explains the lack of SNPs associated with bTB positive samples. In European taurine cattle there is strong evidence for genetic variation in resistance to bTB (Brotherstone et al., 2010; Bermingham et al., 2014; Tsairidou et al., 2014; Woolliams et al., 2008; Tsairidou et al., 2018) with heritabilities of 18% (SE=4) (Bermingham et al., 2009). This has led to the publication of genetic evaluations for resistance to bTB to allow farmers to select breeding sires with greater genetic bTB resistance (Banos et al., 2017). Importantly, these studies have also revealed that bTB resistance is mostly polygenic (Brotherstone et al., 2010; Bermingham et al., 2014; Tsairidou et al., 2014; Woolliams et al., 2008; Tsairidou et al., 2018; Bermingham et al., 2009).

After accounting for age and sex, our GWAS results showed that there was no evidence of a genetic association at the genome-wide significance level. At the suggestive level, there is an association between *M. bovis* infection and SNPs on chromosome 12 (at base pairs 34703645 - 34780062, genome build *Bos taurus* UMD3.1). This region did not overlap with any other regions identified by studies of bTB resistance in cattle eg. (Bermingham et al., 2014; Finlay et al., 2012; Richardson et al., 2016; le Roex et al., 2013; Tsairidou et al., 2018; Amos et al., 2013; Driscoll et al., 2011; Raphaka et al., 2017). However, one of the genes within ±500Kbp of this region was tumor necrosis factor (TNF). TNF is important in macrophage activation as well as cell recruitment to the site of infection (Lin et al., 2007). In humans it is thought that TNF plays a role in the control of tuberculosis, as a study has shown that humans treated with TNF-neutralizing drugs, have increased susceptibility to tuberculosis (Lin et al., 2007).

When the MAF of the SNPs in the Chr12:34703645 - 34780062 region were compared between mixed and Fulani cattle, no difference was observed. Presumably this was due to the reasons described above. Yet, the median MAF of these SNPs is lower in Asian zebu cattle compared to European and African taurine cattle. This difference in MAF could suggest that selection is happening at these SNPs and requires further investigation.

Although our analyses suggest that bTB resistance is partly controlled by cattle genetics, it is possible that the heritability estimate and SNP effects are inflated due to the lack of knowledge of other potential confounders that are usually accounted for in genetic models. For example, we did not include population structure, polygenic additive effects or other systematic environmental effects which may confound the genetic effects. Furthermore, it was not possible to include abattoir in these models due to the small sample sizes in each group. Breed and abattoir may be strongly confounded as the distribution of breeds across abattoirs is not equal. So leaving out ‘abattoir’ from the models has also increased the risk of falsely assigning breed or ‘cluster’ effects to the additive genetic component of this model. More animals are needed to be genotyped before abattoir level effects can be included.

To conclude, there is a need to reconcile the difference between breed phenotype and genotypes of African cattle. We have shown that there is a lack of genetic difference between the apparent reported breeds and that there is an indication of genetic variation in the resistance to bTB however this needs more evidence. Furthermore, we have highlighted the need for better tools to genotype African cattle populations.

Finally, it is important to understand the challenges faced by livestock in specific settings both in terms of pathogens and the environment, in addition to their intended purpose and how they fit into a defined management system. It is only at this point livestock keepers can then make informed breeding choices, not only against resistance to disease but breeding for production traits they require. By doing this it will create a more practical sustainable breed, which is adapt to different circumstances that fit in with the cultural context and local need. Without considering these wider potential impacts, breed improvement strategies can risk the unintended increase in incidence of diseases such as bTB and also profoundly distort the environmental and breed equilibrium, thus massively affect livelihoods.

## ETHICAL STATEMENT

Although this study involved the use of biological tissue from routine post mortem inspection by veterinary public health officials in Cameroon, no experimental work was done on live vertebrates. Tissue sample collection and processing procedures were conducted according to the manual of best practice with an aim of reducing contamination. Research approval at local level was provided by the supervisory of commercial slaughterhouses sanctioned through the Ministry of Livestock, Fisheries and Animal Industries in Cameroon. The research project was also reviewed and approved based on the Animal Scientific Procedures Act of 1986 by the University of Edinburgh Ethical Review Committee (ERC No: OS02-13).

## AUTHOR CONTRIBUTIONS

BB, RK, AM, FNE, ADW conceived and designed the study; RK, FNE, LB, SM, BB: performed the field work; RC analyzed the data; AM, MS, LB, EC, ADW contributed expertise and analysis tools; RC, AM, MS wrote the first draft paper; All authors read and contributed to the final draft of the paper.

## ACKNOWLEDGMENTS

All the authors would like to thank the local abattoir employees and the MINEPIA veterinary staff and delegates, as well as the staff at the Tuberculosis Reference Laboratory, Bamenda and the Laboratory of Emerging Infectious Diseases, University of Buea, without whom this study would not have been possible.

## SUPPLEMENTAL DATA

### DATA AVAILABILITY STATEMENT

The datasets analysed for this study can be found in the Edinburgh DataShare Repository https://doi.org/10.7488/ds/2722.

