## Supplementary Materials for "A putative genomic map for resistance of *Bos indicus* cattle in Cameroon to bovine tuberculosis"

### *Supplementary Material*

#### 1 SUPPLEMENTARY DATA

The datasets supporting the conclusions of this article are available on the Edinburgh DataShare Repository. `https://doi.org/10.7488/ds/2722`

#### 2 SUPPLEMENTARY TABLES AND FIGURES

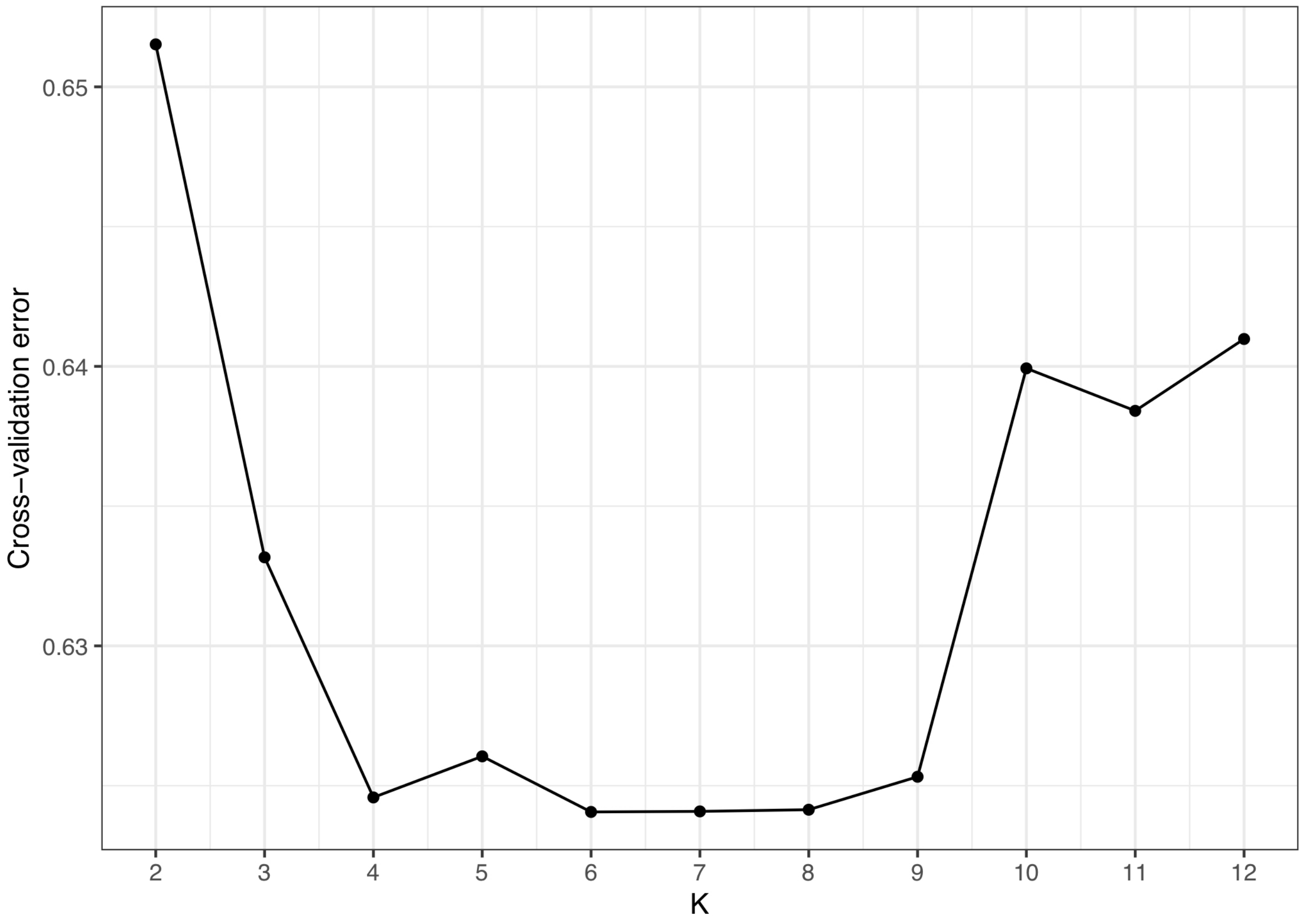


**Figure S1.** Cross-validation plot for the admixture analysis

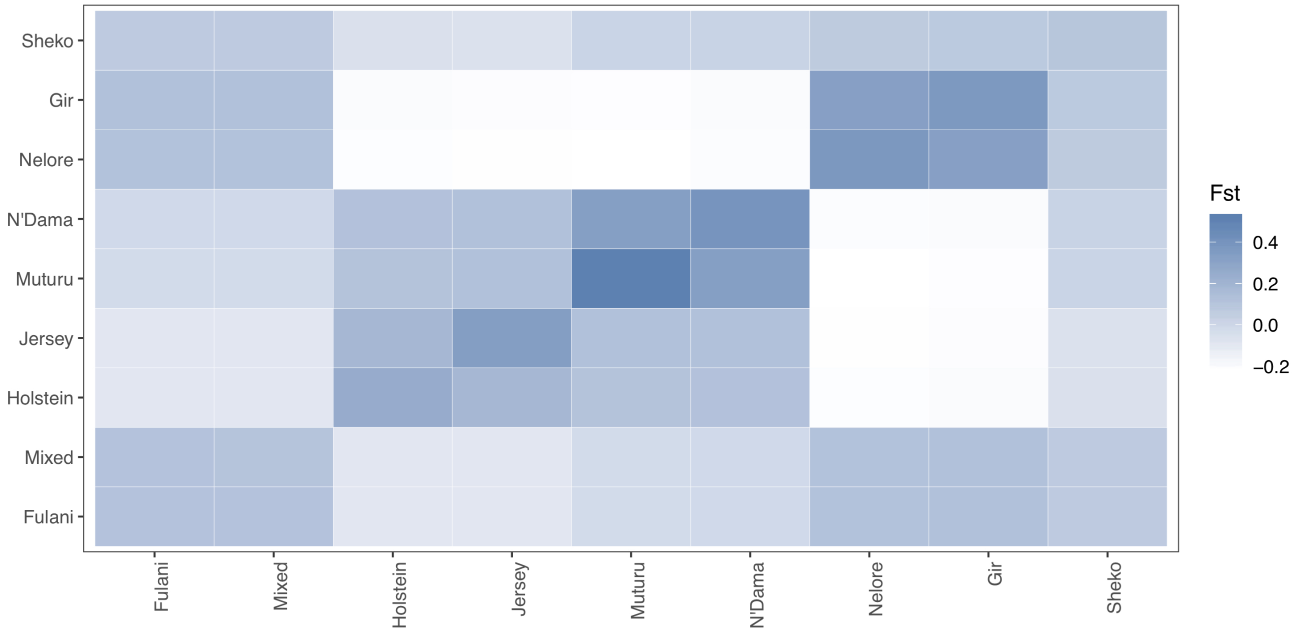


**Figure S2.** Relative beta estimator of Fst using Weir *et al.* 2002 Weir and Hill (2002) method comparing the Fulani and mixed breeds with the reference cattle

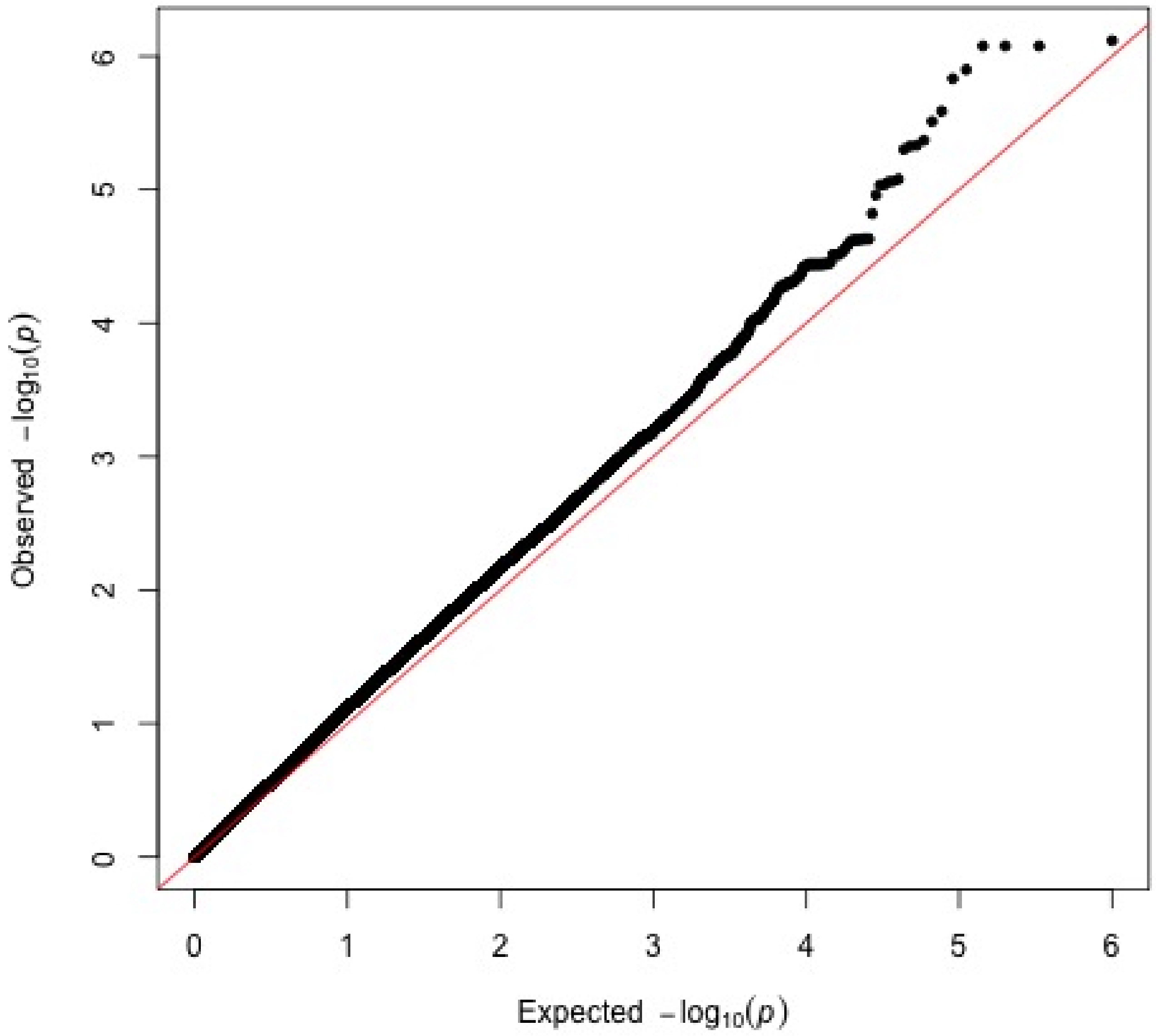


**Figure S3.** QQ-plot for the genome wide association between SNPs and *M. bovis* infection status after accounting for age, sex and breed as covariates.

**Table S1.** The studied cattle populations with number of genotyped individuals in the dataset before ($N_{Before}$) and after ($N_{After}$) quality control. Populations are defined according to DARGIS 2007 DAGRIS (2007).

| Breed | Population | $N_{Before}$ | $N_{After}$ | Source |
|---|---|---|---|---|
| Fulani | Admixed | 172 | 161 | Kelly et al. (2018) |
| Mixed | Admixed | 56 | 52 | Kelly et al. (2018) |
| Holstein | European Taurine | 63 | 17 | Bovine HapMap Consortium (2009) |
| Jersey | European Taurine | 36 | 10 | Bovine HapMap Consortium (2009) |
| Muturu | African Taurine | 10 | 10 | Bahbahani et al. (2017) |
| N'dama | African Taurine | 24 | 21 | Bahbahani et al. (2017) |
| Nelore | Asian Zebu | 35 | 20 | Bahbahani et al. (2017) |
| Gir | Asian Zebu | 30 | 14 | Bovine HapMap Consortium (2009) |
| Sheko | Admixed | 18 | 15 | Bovine HapMap Consortium (2009) |

**Table S2.** The association between bTB lesions and breed using the 164 genotyped animals which had been tested for *Fasciola* sp. using the model structure in Kelly *et al.* 2018 Kelly et al. (2018)

| Predictors | Levels | Odds Ratio | 95% CI | p value |
|---|---|---|---|---|
| Sex | Female | 1.00 | | |
| | Male | 0.32 | 0.13 – 0.80 | 0.014 |
| Age | $\geq$ 3 years | 1.00 | | |
| | $<$ 3 years | 0.71 | 0.25 – 2.00 | 0.512 |
| Breed | Mixed | 1.00 | | |
| | Fulani | 0.28 | 0.10 – 0.74 | 0.010 |
| *Fasciola* sp. | Negative | 1.00 | | |
| | Positive | 2.58 | 0.72 – 9.22 | 0.145 |
| Breed**Fasciola* sp. | | 9.07 | 1.19 – 69.28 | 0.033 |

**Table S3.** The heritability of *M. bovis* infection status. N = number of individuals; Npos = number of positive individuals *M. bovis*; Prevalence = prevalence followed by 95% confidence interval in brackets; VP = phenotypic variance; $h^2$ = heritability. Standard errors are in brackets

| Response Variable | Fixed effects | N | Npos | Prevalence (95% CI) | VP (SE) | $h^2$ (SE) |
|---|---|---|---|---|---|---|
| *M. bovis* infection status | None | 212 | 84 | 39.6 (33.0 - 46.5) | 4.22 (0.722) | 0.218 (0.134) |
| *M. bovis* infection status | Age and sex | 207 | 79 | 38.2 (31.5 - 45.1) | 4.07 (0.836) | 0.189 (0.166) |
| *M. bovis* infection status | Age, sex and breed | 207 | 79 | 38.2 (31.5 - 45.1) | 4.11 (0.84) | 0.198 (0.165) |
| *M. bovis* infection status | Age, sex, breed and abattoir | 207 | 79 | 38.2 (31.5 - 45.1) | 3.96 (1.12) | 0.166 (0.236) |
